# Fibrillarin-1 and Fibrillarin-2 are required for divergent cell lineage development in planarian homeostasis and regeneration

**DOI:** 10.1101/2022.08.08.502111

**Authors:** Jiajia Chen, Xue Pan, Hao Xu, Yuhong Zhang, Kai Lei

**Affiliations:** School of Life Sciences, Zhejiang University, Hangzhou, Zhejiang, China; Westlake Laboratory of Life Sciences and Biomedicine, Key Laboratory of Growth Regulation and Translational Research of Zhejiang Province, School of Life Sciences, Westlake University, Hangzhou, Zhejiang, China; Institute of Biology, Westlake Institute for Advanced Study, Hangzhou, Zhejiang, China

**Keywords:** planarian, ribosome, rRNA modification, epidermal lineage, cell differentiation

## Abstract

Ribosome heterogeneity has been revealed to exist in different cell types during development. However, the function and regulatory mechanisms of ribosome heterogeneity in missing tissue regeneration have yet to be reported. We used the planarian *Schmidtea mediterranea* with whole-body regenerative capability as a model and revealed the function of the rRNA modification protein fibrillarin in cell lineage development and tissue regeneration. We identified two fibrillarin homologs in planarian, *Smed-fbl-1* (*fbl-1*) and *Smed-fbl-2* (*fbl-2*), with distinct expression patterns. While *fbl-2* regulates stem cell proliferation and multiple progenitor cell differentiation, *fbl-1* participates in epidermal lineage late-stage specification and wound response. This study indicates that fibrillarin, a nucleolar protein, can respond to wounds and function in distinct cell types, suggesting the existence and critical roles of ribosome heterogeneity in stem cells and tissue regeneration.

## Introduction

Ribosomal biogenesis is the basis for multiple cellular processes, and its dysfunction influences translation, cell cycle, and immune responses (Bianco & Mohr, 2019; Kang *et al*, 2021; Pelletier *et al*, 2018; Xu *et al*, 2016). Ribosomes are composed of two major components, the large and small ribosomal subunits (LSU and SSU). Each subunit consists of specific ribosomal RNA (rRNA; LSU: 28S rRNA, 5.8S rRNA, and 5S rRNA; SSU: 18S rRNA) and ribosomal proteins (RPs; LSU: RPLs; SSU: RPSs). Ribosome heterogeneity was proposed early in 1958 and revised to the ribosome filter hypothesis in 2002 (Gay *et al*, 2022; Genuth & Barna, 2018; Li & Wang, 2020). Ribosome heterogeneity can be determined by rRNA variants and modifications, RPs, and other factors, including ribosome-associated proteins, subcellular localization of ribosomes, and the ribosomal architecture (Li & Wang, 2020). Its origination remains to be addressed, including distinctly heterogeneous ribosome components, cell-type-specific or subcellular-compartment-specific ribosomes, and recognition and production of specific ribosomes (Gay *et al*., 2022; Genuth & Barna, 2018). rRNA modification is a prominent source of ribosome heterogeneity, with the most abundant type being 2’-O-methylation. Different cell types have diverse profiles of 2’-O-methylation, providing evidence of the ribosome heterogeneity (Krogh *et al*, 2016). Numerous studies have also implied that stem cell differentiation and cell fate determination require increasing ribosome biogenesis and protein synthesis in multiple species (Gay *et al*., 2022; Lv *et al*, 2021; Sanchez *et al*, 2016; Zhang *et al*, 2014). Whether ribosome heterogeneity occurs during stem cell differentiation or cell type specification *in vivo* remains to be explored.

Fibrillarin (FBL) is an essential nucleolar protein that is conserved in organisms ranging from yeast to humans. The main structure of FBL contains two functional domains: a glycine/arginine-rich (GAR) region with nuclear localization signals and an RNA binding domain with methyltransferase (MTase) activity (Pereira-Santana *et al*, 2020; Rodriguez-Corona *et al*, 2015). Depending on its functional domain, FBL catalyzes the 2’-O-methylation of rRNA to process pre-rRNA into 18S rRNA and 28S rRNA. Previous studies have illustrated that FBL functions in the nucleolus to process rRNA through liquid-liquid phase separation via its GAR domain (Yao *et al*, 2019). Guided by small nucleolar RNA (snoRNA), FBL participates in different cellular processes (Li *et al*, 2018; Ren *et al*, 2019; Yi *et al*, 2021). FBL can regulate stem cell viability and control stem cell differentiation through the p53 signaling pathway (Watanabe-Susaki *et al*, 2014). Inhibition or mutation of FBL influences the mouse early development (Newton *et al*, 2003), arrests cells at the S-phase of the cell cycle and impairs neuron differentiation in zebrafish (Bouffard *et al*, 2018). Considering the role of FBL in stem cell biology, we hypothesized that FBL generates ribosome heterogeneity by rRNA modification to regulate the development of stem cells during tissue homeostasis and regeneration.

Planarian is considered as a remarkable animal model that can be used to investigate regeneration mechanisms. Using planarians, which have the whole-body regenerative capability, allowed us to explore the functions of genes from the perspective of the stem cell biology (Reddien, 2018). Proper neoblast proliferation and differentiation are critical for cell renewal and regeneration after amputation or injury (Scimone *et al*, 2014). The precise regulation of neoblasts self-renew and differentiate to replace missing tissue or maintain homeostasis is still being studied (Bohr *et al*, 2021; Boser *et al*, 2013; Lei *et al*, 2016; Reddien, 2018, 2021; Wagner *et al*, 2012; Wenemoser & Reddien, 2010).

We found two homologs of FBL in planarians. Regenerative capacity was destroyed after the knockdown of *Smed-fbl-1* (*fbl-1*) or *Smed-fbl-2* (*fbl-2*). In line with the function of FBL, the rRNA expression levels were decreased in animals subjected to RNA interference (RNAi). Evidence regarding cell-type-specific ribosomes was obtained via analysis of the cell types differentially expressing *fbl-1* and *fbl-2. fbl-1* was explicitly expressed in *AGAT-1*^+^ cells and their mature descendants, whereas *fbl-2* was expressed in progenitor cells and stem cells. Upon dissecting the functions of the two homologs in planarians during homeostasis and regeneration, we propose that *fbl-1* and *fbl-2* differentially regulate cell proliferation, differentiation, and even epidermal cell specification through specific ribosome translation.

## Results

### Knockdown of *Smed-fbl-1* or *Smed-fbl-2* shows distinct defects in homeostasis

The human FBL sequence was used to search for homologs in the planarian *Schmidtea mediterranea* by protein sequence alignment. Both evolutionary conservation and cellular expression patterns validated the homologs of FBL in planarians. SMED30025294 and SMED30005569, termed *fbl-1* and *fbl-2*, respectively, were predicted to encode FBL after performing a phylogenetic comparison and analyzing the GAR and MTase functional domains (Fig 1A). Distinct from the nucleolar localization of both FBL proteins in H9 cells, FBL-2 is located at the cytoplasm in 293T cells, suggesting the different cellular localization of FBL-2 in various cell types (Fig EV1A and B).

**Figure 1.**
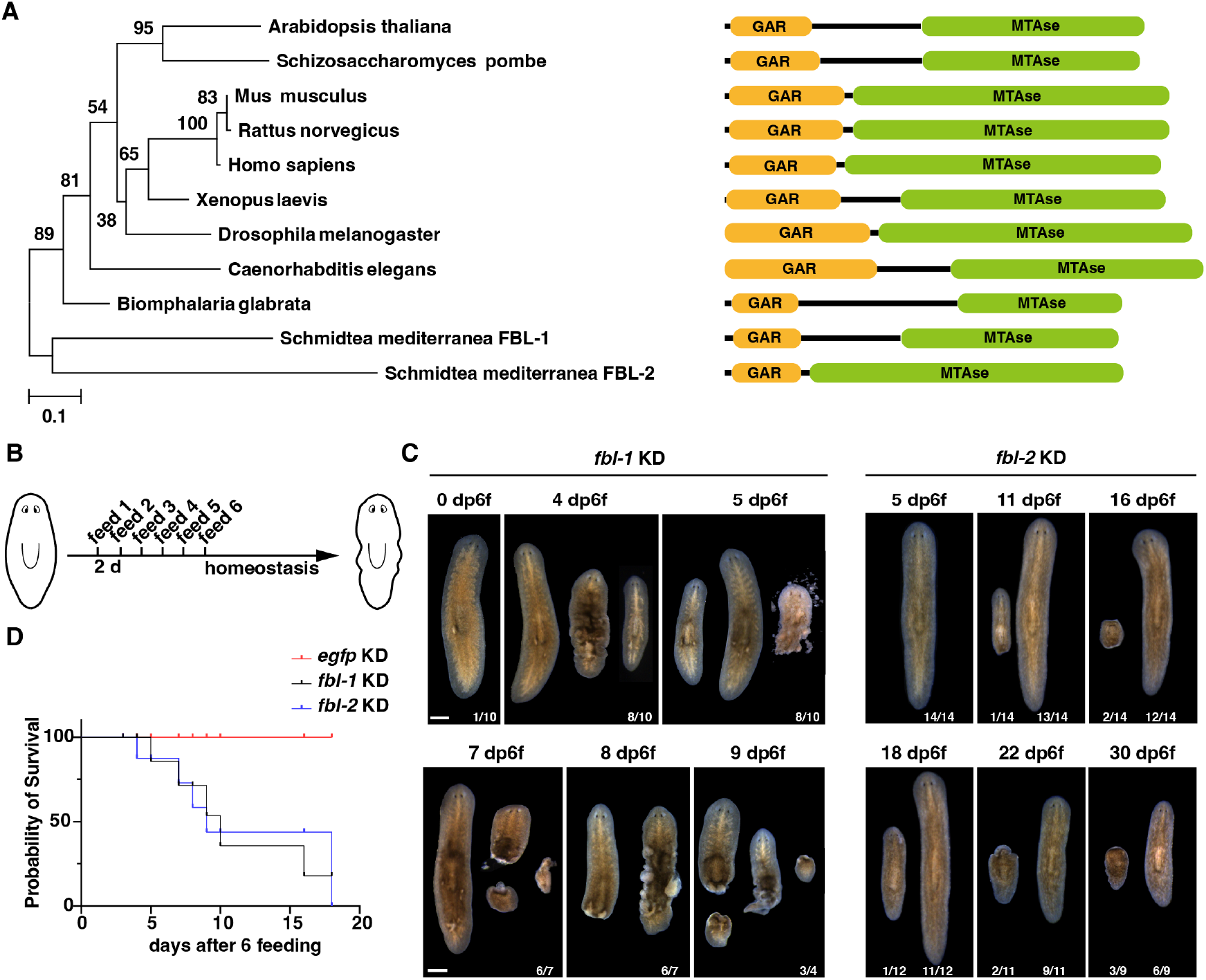
*fbl-1* and *fbl-2* are required for tissue homeostasis. A. Phylogenetic tree of FBL protein homologs from 10 species with a schematic diagram of the functional domains. The posterior probability value is shown on each branch. B. Feeding schedule (six times every three days) to examine the function of *fbl-1* and *fbl-2* in homeostasis. C. Different effects on homeostasis caused by KD of *fbl-1* or *fbl-2*. White spots on the dorsal surface and regressing heads of *fbl-2* KD worms in a small population during homeostasis were observed. The number indicates the penetration of defective phenotypes in representative images. Scale bars, 500 μm. D. Survival curves of animals after KD of *fbl-1* or *fbl-2*.

To investigate the functions of these homologs in planarians, we performed RNAi experiments to knock down the expression of *fbl-1* and *fbl-2*, respectively. Both *fbl-1* and *fbl-2* are required for homeostasis, supporting the essential function of FBL at the same feeding schedule (Figs 1B-D, EV1C). Upon observation of the defects, *fbl-1* knockdown (KD) animals showed white spots on the dorsal epidermis before tail breaking and body disassembly (lethal phenotype) (Fig 1C). By contrast, 57% of *fbl-2* KD animals displayed head regression and then disassembly, while others remained intact (Fig 1C). These phenomena suggested that *fbl-1* and *fbl-2* play diverse roles in homeostasis.

### *Smed-fbl-1* and *Smed-fbl-2* are required for homeostasis and regeneration by modulating rRNA expression

To observe the regenerative capacity of *fbl-1* KD animals, we modified the RNAi feeding schedule as shown in Figure 2A. The animals still displayed different effects on homeostasis after KD of *fbl-1* or *fbl-2* (Fig 2B and C). Lesions appeared in the tail of *fbl-1* KD animals, but *fbl-2* KD animals exhibited head regression (Fig 2B and C). These phenotypes led us to explore the functional difference between *fbl-1* and *fbl-2* in planarians. Quantitative real-time PCR (qPCR) was used to confirm the KD efficiency. The results showed that KD of *fbl-1* did not influence the expression of *fbl-2*, and vice versa (Fig 2D and E). Furthermore, the decreased expression of FBL-2 after KD of *fbl-2* was validated by anti-FBL-2 antibody immunofluorescent staining in *fbl-2* KD animals compared with that in *egfp* KD controls (Fig EV1D). To confirm the functions of *fbl-1* and *fbl-2* in rRNA biogenesis, we performed qPCR to measure the relative amounts of 18S rRNA and 28S rRNA in *fbl-1* or *fbl-2* KD planarians, respectively. Both *fbl-1* KD and *fbl-2* KD planarians had significantly lower 18S rRNA and 28S rRNA levels than *egfp* KD controls (Fig 2F-I). Taken together, these results suggested that *fbl-1* and *fbl-2* are conserved to modulate rRNA biogenesis.

**Figure 2.**
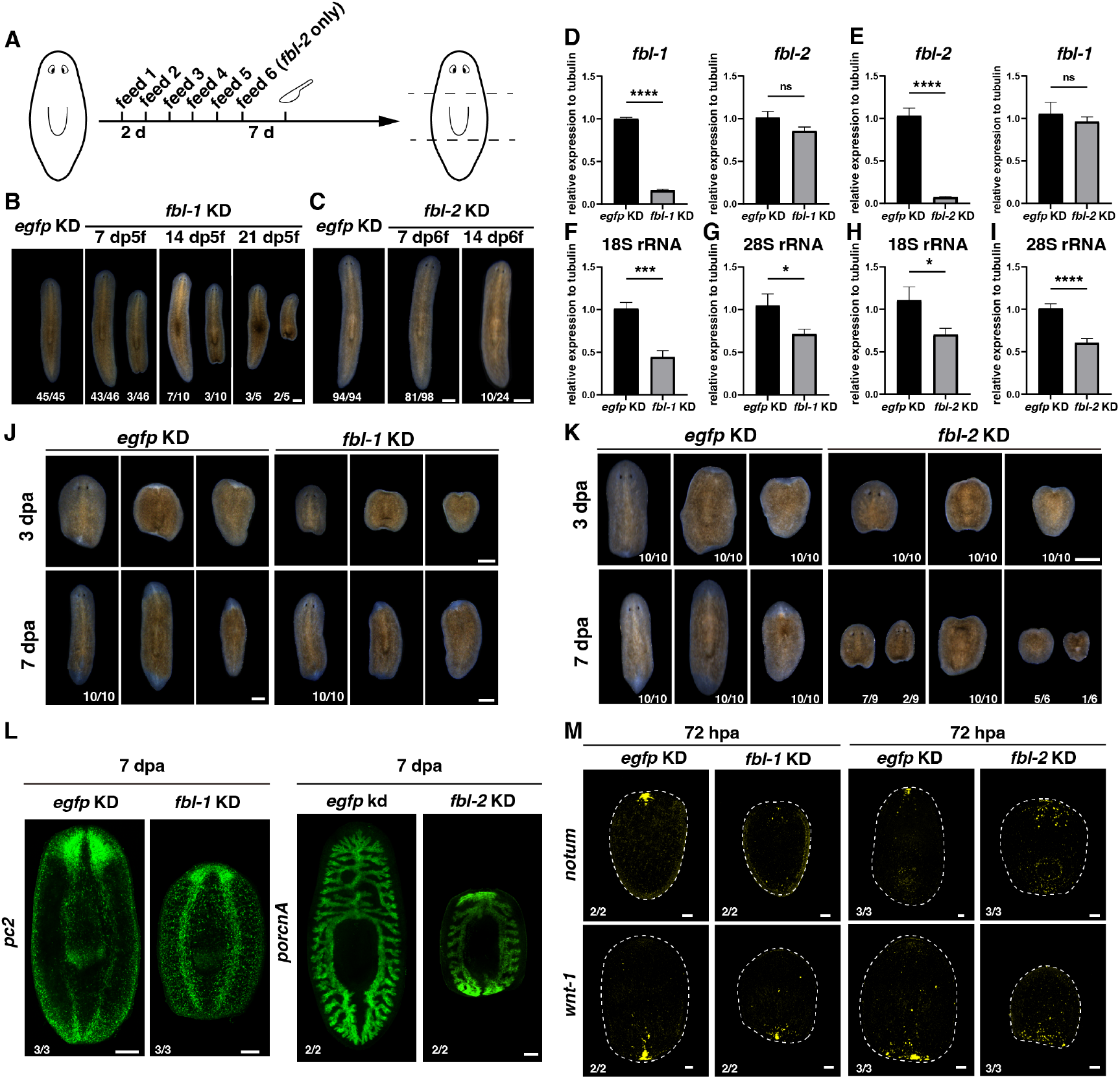
*fbl-1* and *fbl-2* are required for tissue homeostasis and regeneration. A. Schedule of RNAi feeding every three days and amputation of *fbl-1* and *fbl-2* KD animals. B, C. Different effects of *fbl-1* KD (B) and *fbl-2* KD (C) on homeostasis under a modified feeding schedule. Scale bars, 500 μm. D. Relative mRNA level of *fbl-1* and *fbl-2* in *fbl-1* KD animals measured by qPCR. E. Relative mRNA level of *fbl-1* and *fbl-2* in *fbl-2* KD animals measured by qPCR. F, G. Relative expression levels of 18S rRNA and 28S rRNA after *fbl-1* KD. H, I. Relative expression levels of 18S rRNA and 28S rRNA after *fbl-2* KD. ns, not significant; \**P* < 0.05; \*\*\**P* < 0.001; \*\*\*\**P* < 0.0001 (Student’s *t*-test). *J. fbl-1* KD causes regenerative delay. Scale bars, 500 μm. K. Regenerative defects of *fbl-2* KD animals at 3 and 7 dpa. Scale bar, 500 μm. L. Regeneration effects are shown by mature tissue marker expression at 7 dpa (neurons, *pc2*; gut, *porcn A*). Scale bars, 200 μm. M. Abnormal expression of anterior and posterior pole markers (*notum, wnt-1*) at 72 hpa. Scale bars, 100 μm.

To examine the functions of *fbl-1* and *fbl-2* in missing tissue regeneration, we amputated the KD animals seven days after the last feeding. The *fbl-1* KD animals regenerated more slowly and exhibited smaller blastema, while the *fbl-2* KD animals could not regenerate the anterior and posterior tissues (Figs 2J-K, EV2A). The blastema formation involves the expression of wound response genes, proliferation and differentiation of neoblasts, and guidance of the positional signals (Reddien, 2018; Wenemoser *et al*, 2012; Wenemoser & Reddien, 2010). However, a comparison of the levels of early response genes (*fos1, jun1*, and *runt1*) showed normal induction of the wound response in *fbl-2* KD animals (Fig EV2B). The staining of various tissues confirmed the regeneration defects after KD of *fbl-1* or *fbl-2*, which showed incompletely regenerative CNS and intestinal branches at 7 days post-amputation (dpa) (Fig 2L). The regeneration defects were further confirmed by the disrupted formation of anterior (*notum*) and posterior (*wnt-1*) polarity in both *fbl-1* KD and *fbl-2* KD animals at 72 hours post-amputation (hpa) (Fig 2M). Unlike *fbl-1* KD animals, the expression of *notum* and *wnt-1* is obviously affected in *fbl-2* KD animals, suggesting the disturbed expression of positional control genes by *fbl-2* KD.

### *Smed-fbl-1* and *Smed-fbl-2* are expressed in distinct cell types

To examine how *fbl* inhibition causes regeneration and homeostasis defects, whole-mount in situ hybridization (WISH) and fluorescent in situ hybridization (FISH) of planarians were performed to elucidate the expression patterns of *fbl-1* and *fbl-2* (Fig 3A, F). The spatial expression pattern along the DV axis showed that the *fbl-1*^+^ cells were distributed from the dorsal mesenchyme to the dorsal epidermis, but the *fbl-2*^+^ cells were distributed extensively (Fig 3A, F). Based on the above expression patterns, we first proposed that *fbl-1* is expressed in epithelial lineage cells. The development of planarian epithelial cells has been identified with multiple progenitor types through the lineage (Cheng *et al*, 2018; Tu *et al*, 2015; Wurtzel *et al*, 2015; Zhu *et al*, 2015; Zhu & Pearson, 2018). The *prog-1*^+^ and *AGAT-1*^+^ cells have been identified as early and late progenitors of the epithelial lineage (Eisenhoffer *et al*, 2008). *AGAT-1*^+^ cells have been further characterized as specifiers of mature epithelium, while *vim-3*^+^ cells are produced after the *zpuf-6*^+^ transient state (Tu *et al*., 2015). Therefore, through double FISH (dFISH) to co-label *fbl-1*^*+*^ cells with epithelial lineage markers, we showed that *fbl-1* is expressed in epithelial lineage cells, distinctly in *AGAT-1*^+^ cells. The ratio of cells co-expressing *AGAT-1* and *fbl-1* was high (agat-1/fbl-1: 99.5%), whereas smaller proportion of *fbl-1* co-expression was observed in *vim-3*^+^ or *zpuf-6*^+^ cells (zpuf-6/fbl-1: 11.6%, vim-3/fbl-1: 31.9%) (Fig 3B). Furthermore, *fbl-1* transcript levels were increased after amputation at the anterior-faced or posterior-faced wound sites during regeneration initiation (4-24 hpa), suggesting a wound response at the level of *fbl-1* transcription (Fig 3C). Notably, the expression of *fbl-1* retained a pattern similar to that in intact worms and was comparable to that in the control group at 48 hpa (Figs 3C, EV3C). At 72 hpa, the knockdown efficiency was restored, and *fbl-1* was highly co-expressed within *vim-3*^+^ cells (Fig 3D and E). This evidence suggested that ribosome biogenesis and protein synthesis are required during stem cell differentiation, which triggers high expression of *fbl-1* (overload) rather than inhibiting *fbl-1*. In addition, *fbl-1* was not expressed at the blastema at 48 hpa, where *AGAT-1*^+^ cells will accumulate, suggesting that *fbl-1* is expressed in the late stage of *AGAT-1*^+^ cells without affecting their generation.

**Figure 3.**
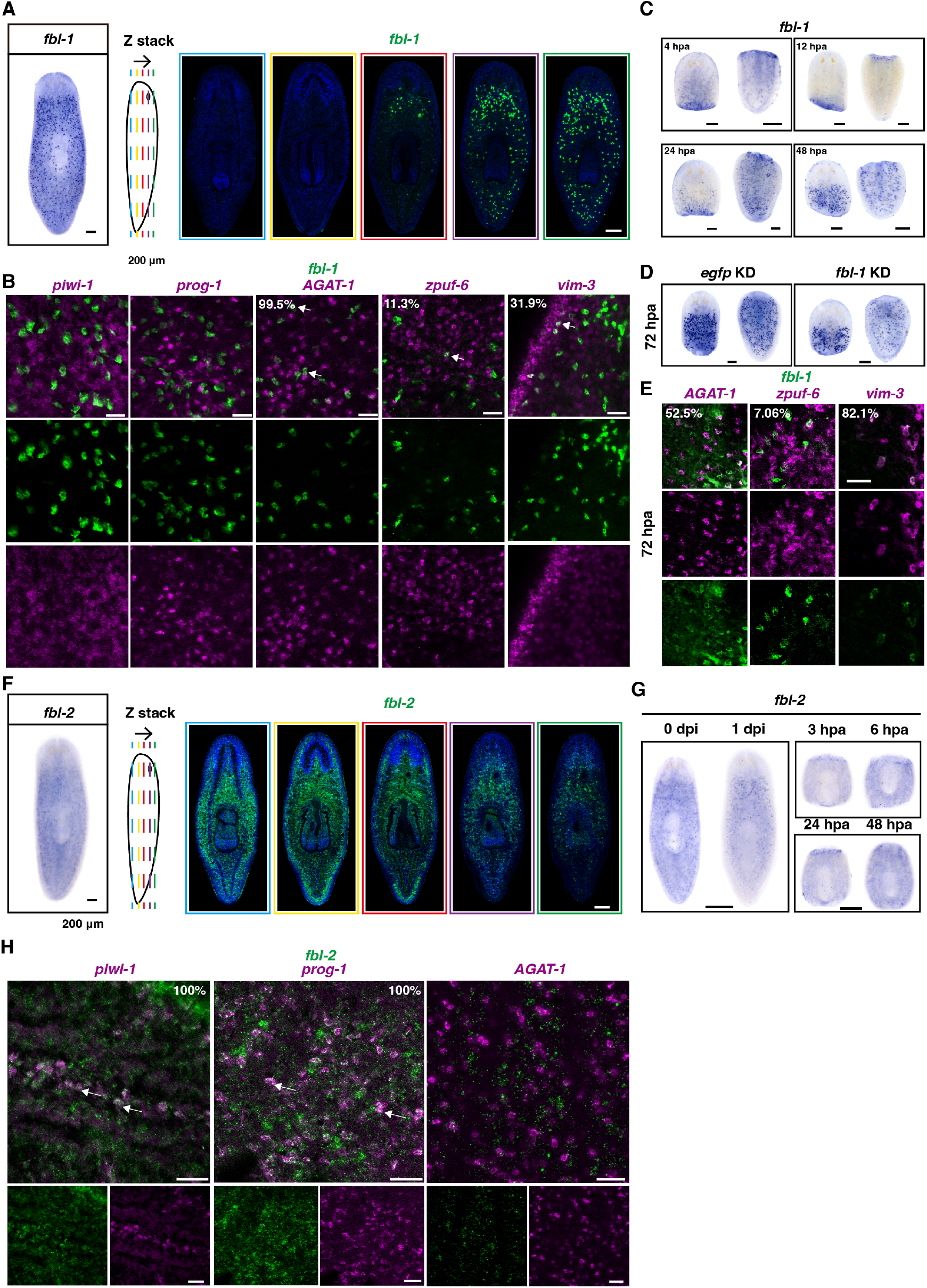
Expression analysis of *fbl-1* and *fbl-2* in distinct cell lineages. A. The expressed pattern of *fbl-1* in planarians by FISH and *fbl-1* expression pattern is shown as the z stack in planarians. Scale bars, 200 μm. B. Colocalization of *fbl-1* with epithelial lineage markers (*prog-1, AGAT-1, zpuf-6*, and *vim-3*) in intact worms. The percentages of marker positive cells to *fbl-1* positive cells are shown. Scale bars, 500 μm. C. *fbl-1* expression responds to injury at the wound site from 4 hpa to 24 hpa. Scale bars, 200 μm. D. No distinct decrease in *fbl-1* expression was observed at 72 hpa compared with the expression in the control group. Scale bars, 500 μm. E. Colocalization of *fbl-1* with epithelial lineage markers (*AGAT-1, zpuf-6*, and *vim-3*) at 72 hpa. The percentages of marker positive cells to *fbl-1*^+^ cells are shown. Scale bars, 100 μm. F. The expression pattern of *fbl-2* in planarians as determined by WISH and the *fbl-2* expression pattern shown as a z stack. Scale bars, 200 μm. G. *fbl-2* expression levels at 0 and 1 dpi and at 3, 6, 24, and 48 hpa. Scale bars, 500 μm. H. Colocalization of *fbl-2* with a stem cell marker (*piwi-1*), epithelial progenitor marker (*prog-1*), and mature epithelial cell marker (*AGAT-1*). The percentages of *fbl-2* positive cells with indicated cell type markers are shown. White arrow: double-positive cells. Scale bars, 50 μm.

We second predicted that *fbl-2* is expressed in stem cells or progenitor cells based on the evidence that 1) KD of *fbl-2* caused head regression, reflecting a defect in stem cells (Reddien *et al*, 2005); 2) most *fbl-2*^+^ cells were depleted at one-day post-irradiation (dpi) (Fig 3G). The dFISH experiments showed that *fbl-2* was expressed widely in different cell types. Given that *fbl-1* was expressed in epidermal lineage, we next examined whether *fbl-2* also facilitated epidermal cell differentiation. The signals of *fbl-2* co-expressed in *prog-1*^+^ cells were evident, whereas those of *fbl-2* co-expressed in neoblasts were dot-like (Figs 3H, EV3E). Some unknown cell types also expressed *fbl-2* (Fig EV3A and B). In addition, *fbl-2* was stimulated at the wound site at 24 hpa when the stem cells were activated for proliferation and differentiation (Fig 3G). This finding is consistent with the co-expression of *fbl-2* with early progenitor and stem cell markers. In contrast to the expression of *fbl-1* at the early time point after amputation, *fbl-2* did not respond to wounds but was parallel to the waves of H3P expression at 6 and 48 hpa, which is also consistent with the normal expression of wound-induced genes after KD of *fbl-2* (Figs EV2B, EV3D). The above results suggested that FBL-1 functions in epidermal cell lineage development, especially the specification from *AGAT-1*^*+*^ cells to *zpuf-6*^+^ and *vim-3*^+^ cells, while FBL-2 functions in stem cells and progenitor cells to regulate cell proliferation or differentiation.

### *Smed-fbl-2* is required for stem cell proliferation and differentiation into early progenitors

To elucidate the cellular mechanisms regulated by *fbl-1* and *fbl-2*, we first examined the stem cell population (*piwi-1*^+^) and proliferating cell population (H3P^+^) in *fbl-1* and *fbl-2* KD worms. Compared with *egfp* KD and *fbl-1* KD worms, only *fbl-2* KD planarians exhibited reductions in the number of proliferative cells during homeostasis, consistent with the expression of *fbl-2* in neoblasts and the head regression phenotype (Figs 4A and B, EV5A-D). The *fbl-2* KD animals almost maintained complete but shrank bodies. The number of neoblasts (*piwi-1+*) decreased slightly at seven days post-feeding (dpf) (Fig EV4A). However, the H3P signals were significantly reduced after KD of *fbl-2* during homeostasis and regeneration (Fig 4B and C). It is possible that some stem cells cannot enter the M phase of the cell cycle after *fbl-2* inhibition, similar to the previous discovery that *fbl* promotes the cell cycle in zebrafish (Bouffard *et al*., 2018).

**Figure 4.**
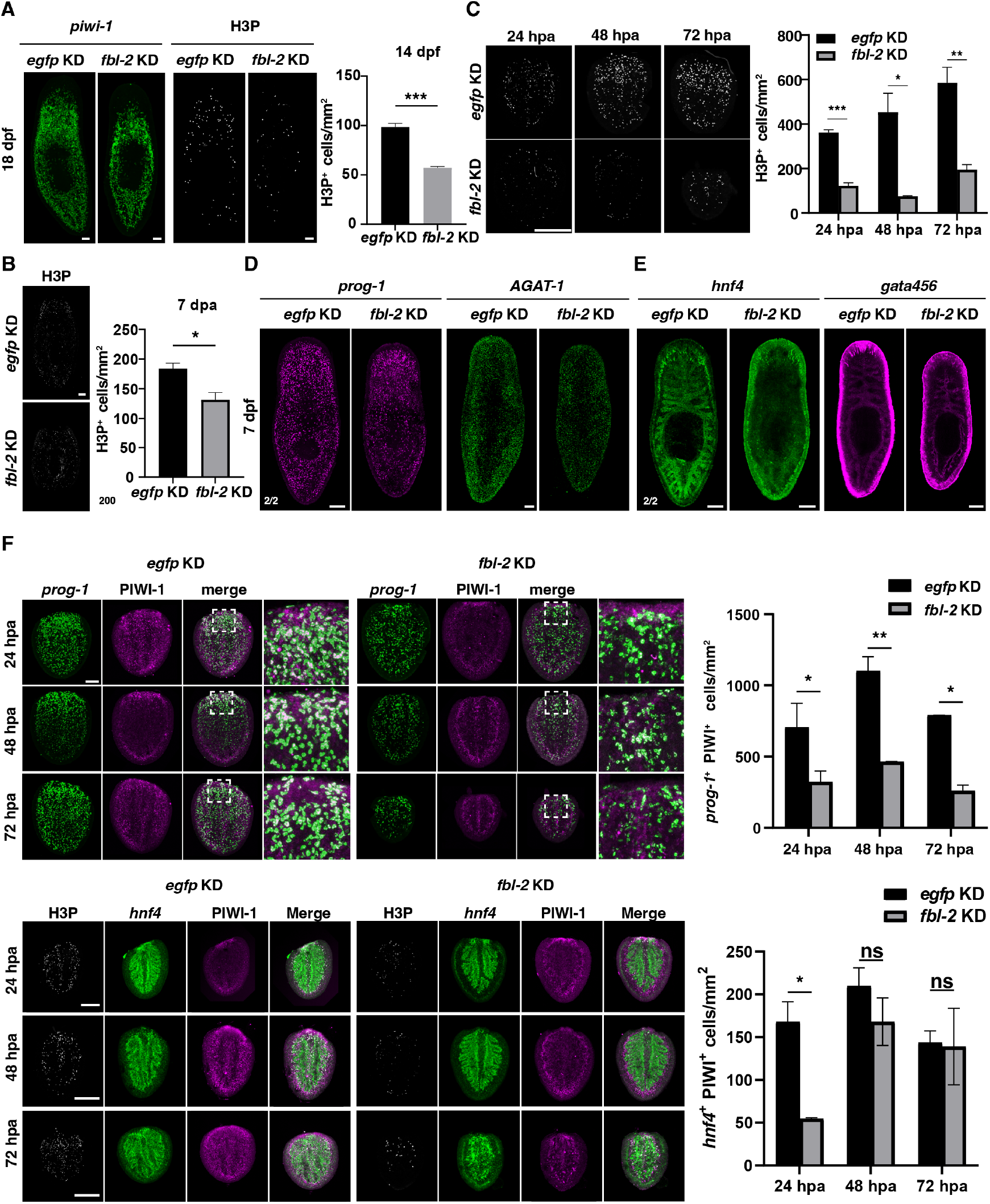
Reduction in cell proliferation and disruption of multiple cell lineage differentiation after *fbl-2* KD. *A*. There was no significant difference in stem cells during homeostasis, but a decreased number of proliferated cells, as shown by H3P signals. Scale bars, 100 μm. \*\*\**P* < 0.001 (Student’s *t*-test). *B*. Cell proliferation and quantification of H3P during regeneration. Scale bar, 200 μm. \**P* < 0.05 (Student’s *t*-test). *C*. Cell proliferation and quantification of H3P^+^ cells after *fbl-2* KD during regeneration. Scale bar, 500 μm. *D*. FISH images show the expression levels of the epidermal progenitor marker *prog-1* and the maturity marker *AGAT-1* during homeostasis in *egfp* KD and *fbl-2* KD animals at 7 dpf. Scale bars, 200 μm. *E*. FISH images show the expression levels of the intestinal progenitor marker *hnf4* and the maturity marker *gata456* during homeostasis in *egfp* KD and *fbl-2* KD animals at 7 dpf. Scale bar, 200 μm. *F. fbl-2* KD blocks progeny (*prog-1*^*+*^, *hnf4*^*+*^) differentiation at 24, 48, and 72 hpa, as shown by FISH and quantitation of double positive cells with PIWI-1. Scale bars, 200 μm. \**P* < 0.05; \*\**P* < 0.01 (Student’s *t*-test).

Considering the *fbl-2* expression in progenitor cells, we suspected that *fbl-2* was also associated with cell differentiation to maintain tissue homeostasis. Therefore, *fbl-2* KD worms at 7 dpf were chosen to identify the expression levels of progenitors. Notably, the expression of *prog-1* (an epidermal early progenitor marker) and *hnf4* (an intestinal progenitor and mature cell marker) was disturbed in *fbl-2* KD animals (Fig 4D and E), which further caused abnormalities in epidermal (*AGAT-1*^+^) and intestine (*gata456*^+^) morphology. Moreover, the newly differentiated progenitors decreased within the blastema in *fbl-2* KD animals, as indicated by the reduction of *prog-1*^*+*^PIWI-1^*+*^, *hnf4*^*+*^PIWI-1^*+*^, and *ovo*^*+*^PIWI-1^*+*^ cells compared with those in *egfp* KD controls at 24, 48, and 72 hpa (Figs 4F, EV4B). In conclusion, KD of *fbl-2* blocked stem cell proliferation and differentiation during homeostasis and regeneration.

### *Smed-fbl-1* is required for epidermal integrity and wound response

Given the epithelial expression pattern and lesion in tails, we assessed the mature tissue markers to narrow down the specific defects induced by KD of *fbl-1*. The staining of muscle, epidermis, and neurons indicated that *fbl-1* KD destroyed multiple tissue morphologies in homeostasis, which might be explained by a secondary effect of epidermal disruption (Fig 5A). Remarkably, *fbl-1* KD planarians displayed an abnormal pattern of epidermal cells distinct from that of *fbl-2* KD planarians (Fig 5B). Therefore, we chose different time points to examine the dynamic expression of epithelial cells. The distribution of *AGAT-1*^+^ cells in planarians can be separated approximately into three layers according to the morphology and number along the LM axis. FISH and quantification analysis indicated that the total number of *AGAT-1*^+^ cells decreased, especially those cells closed to the body boundary (layer 1) at 7 dpf (Fig 5C and D). Until 21 dpf, all the worms exhibited damaged tails or died, the expression of *vim-3* in the wound site was disturbed, and *AGAT-1*^+^ cells were accumulated (Fig 5E). Consistent with the defects in homeostasis, loss of epithelial integrity and cell numbers also occurred to *fbl-1* KD worms after amputation (Figs 5F and G, EV5G). These results suggested that during homeostasis and regeneration, *fbl-1* is required for the specification of *AGAT-1*^+^ cells into the mature epidermis (e.g., *vim-3*^+^ cells).

**Figure 5.**
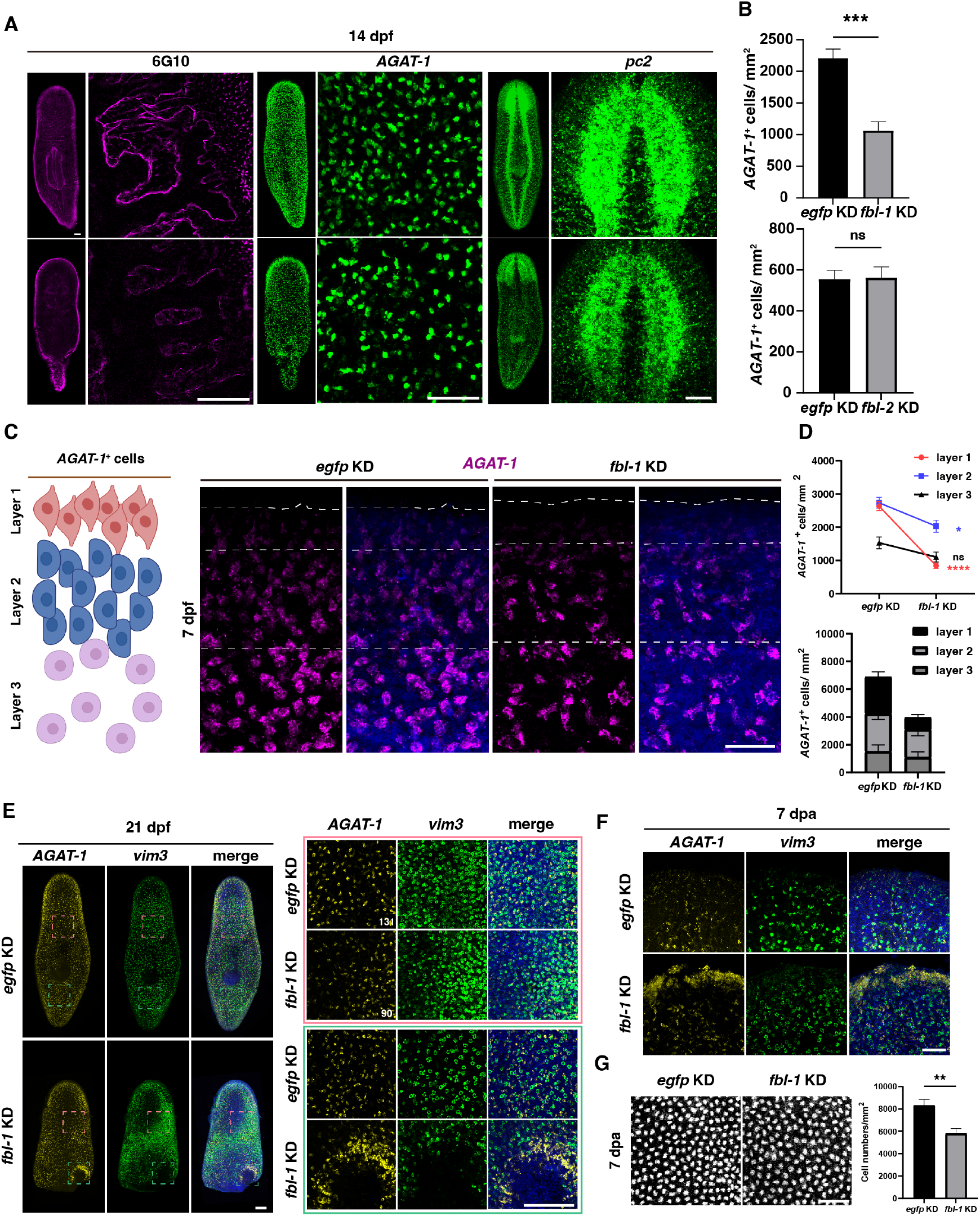
*fbl-1* facilitates the epidermal specification and serves as a wound-induced gene to affect proliferation and cell death during regeneration. *A. fbl-1* KD causes the loss of muscle, epithelial, and neuronal tissue during homeostasis. Scale bars, 100 μm. *B*. Quantification of *AGAT-1*^+^ cells in *fbl-1* KD and *fbl-2* KD animals at 14 dpf. \*\*\**P* < 0.001 (Student’s *t*-test). *C*. Disturbed *AGAT-1*^+^ cells from FISH-labeled *AGAT-1*^+^ cells far from the boundary of the body wall after *fbl-1* KD. Scale bar, 50 μm. *D*. Quantification of *AGAT-1*^+^ cells in three layers of the animal body between the *fbl-1* and *egfp* KD groups at 7 dpf. ns, not significant; \**P* < 0.05; \*\*\*\**P* < 0.0001 (Student’s *t*-test). *E*. Loss of cells positive for the epidermal lineage marker *vim-3* in broken tails during homeostasis 21 dpf. Scale bars, 200 μm. *F. AGAT-1*^+^ cells are stimulated at the wound site at 7 dpa after *fbl-1* KD. Scale bar, 100 μm. *G*. DAPI staining and quantification show decreased epidermal cell number after *fbl-1* KD at 7 dpa. Scale bar, 50 μm.

Amputation triggers wound response and neoblasts proliferation to restore missing tissue (Wenemoser & Reddien, 2010). The increase of *fbl-1*^+^ cells at wound sites and delayed regeneration after KD of *fbl-1* prompted us to explore the balance between cell proliferation and cell death during regeneration. There are two mitotic peaks at 6 and 48 hpa, whereas cell death is increased at 4 and 72 hpa. We found that KD of *fbl-1* would cause a significant reduction of H3P^+^ cells at 48 hpa and increased TUNEL^+^ signals at 72 hpa (Fig EV5E and F). Combined with the results of the analysis above, this evidence indicated that *fbl-1* is required for wound responses and epidermal integrity.

In summary, we proposed that two genes encoding homologs of FBL are essential for planarian tissue homeostasis and regeneration in distinct manners (Fig 6A and B). The *fbl-1* was expressed in epidermal lineage cells and regulated the specification of *AGAT-1*^+^ cells. The reduced regenerative ability resulted from the loss of wound response at the early time points and from decreases in protein synthesis to trigger epidermal maturation after KD of *fbl-1*. Unlike *fbl-1, fbl-2* helped maintain the proliferation of stem cells and regulate the differentiation of multiple cell-lineage progenitors (Fig 6A). The specific modification of FBL in distinct cell types increased the ribosome heterogeneity to preferentially translate the protein required for particular cell types (Fig 6B).

**Figure 6.**
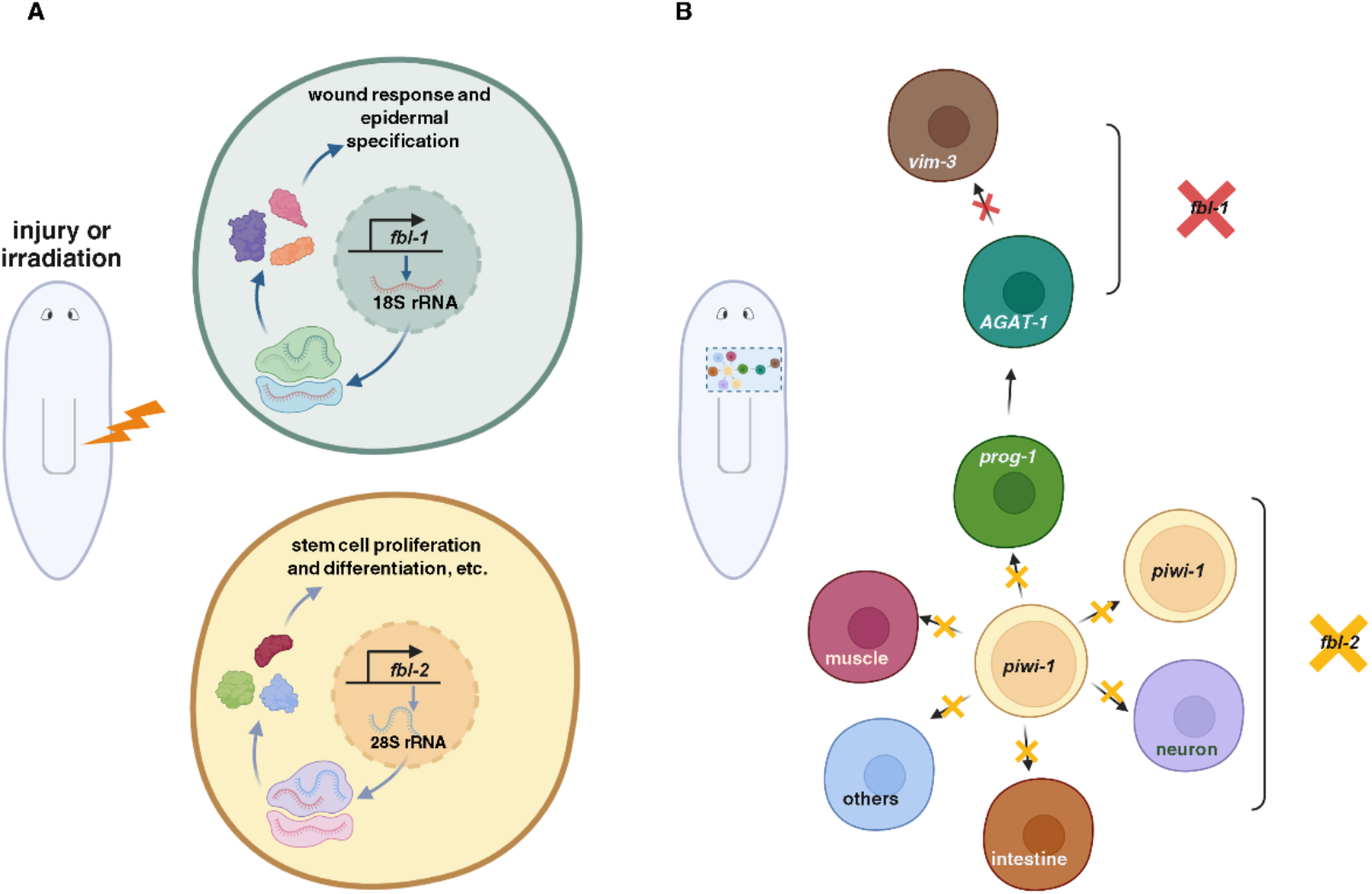
Working model suggests the existence of ribosome heterogeneity regulating cell regeneration during homeostasis and in the context of missing tissue regeneration. *A. fbl-1* and *fbl-2* regulate planarian homeostasis and regeneration via ribosome heterogeneity. *B. fbl-1* and *fbl-2* inhibition block the cell fate transition.

## Discussion

FBL was first identified as a nucleolar protein in 1958 (Crick, 1958). This protein is conserved in organisms ranging from archaea to humans and catalyzes rRNA 2’-O-methylation and pre-rRNA processing via phase separation (Yao *et al*., 2019). FBL is a useful nucleolar marker to examine the dynamics or morphology of the nucleolus during multiple cellular processes. However, numerous studies have demonstrated its functions in regulating stem cell survival, proliferation, differentiation, and even embryonic development in mice, which explicitly utilize rRNA modification and rRNA transcription (Morral *et al*, 2020; Zhang *et al*., 2014). Recent studies have proven the existence of ribosome heterogeneity in Drosophila, which can be decided by rRNA variants and RPs (Genuth & Barna, 2018). The KD of *fbl* will reduce the rRNA modification, which can be used as a tool to elucidate how ribosome heterogeneity determines cell fate and nucleolar protein-regulated tissue regeneration (Erales *et al*, 2017; Jansson *et al*, 2021). Therefore, we investigated FBL in distinct planarian cell types to link its function with cell lineage development underlying ribosome heterogeneity (Fig 6A and B). This provides evidence to understand how epidermal cell differentiation is precisely regulated by the differential expression of *fbl-1* and *fbl-2* in early and late stages, which is divergent from other cell lineages such as intestinal and neural lineages. Further studies on how FBL-mediated modification of rRNA sites and the recognition of mRNA by cell-type-specific ribosomes regulate the “on” and “off” cell fate decisions are needed.

Regeneration requires a precise response to missing tissue. Previous studies on wound response gene expression have revealed that three cell types are activated in response to wounding, including neoblasts, epidermal cells, and muscle cells, in which cell-type-specific genes are expressed (Wurtzel *et al*., 2015). In parallel with these findings, *fbl-1*^+^ epidermal cells responded to wounds, and the regenerative capacity was impaired after *fbl-1* inhibition. Given the multiple distinct steps of the epidermal lineage development, this discovery may explain how ribosome heterogeneity during regeneration regulates cell-type-specific gene expression in epidermal cells (Pearson & A, 2010; Scimone *et al*, 2010; Tu *et al*., 2015; van Wolfswinkel *et al*, 2014; Zhu *et al*., 2015; Zhu & Pearson, 2018). Adopting an additional homolog of FBL provides an advantage in efficiently regulating the sequential requirement of ribosome heterogeneity for further differentiation beyond *AGAT-1*^+^ cells. Although regeneration capacity was diverse among multiple species, the depletion of *fbl* destroyed the transition of stem cells states in mice and zebrafish (Bouffard *et al*., 2018; Wu *et al*, 2022). All these findings suggested that the ability of FBL to regulate stem cell differentiation in physiological homeostatic and tissue regeneration may be conserved among multiple species.

## Materials and methods

### Animal maintenance and irradiation

The *Schmidtea mediterranea* CIW4 planarian strain was cultured in 1X Montjuic water at 20°C. The experimental animals were starved for up to 7 days. For irradiation, 4 ∼ 5 mm worms were exposed to 60 Gy of radiation by an RS2000 Pro X-ray irradiation apparatus.

### RNA extraction, gene cloning, and expression analysis

Total RNA was extracted with TRIzol (Invitrogen, 10296010) and reverse-transcribed into cDNA with ABScript II-RT Mix for qPCR with gDNA Remover (ABclonal, RK20403) for gene cloning. The primers for gene cloning were designed according to the transcriptome database (https://planosphere.stowers.org/), and the primers for qPCR were designed by an online website (https://simrbase.stowers.org/cgi-bin/primer3.pl), as follows: *fbl-1*: 5’ AAGGCCTCTTGTATCGATTC and 3’ CCATTGACAGCCAAGATTTC; *fbl-2*: 5’ TTTCCAGCATCAGGTGAAAG and 3’ ATCTCCAAATCCTCCCCTAC; 18S rRNA: 5’ AACGGCTACCACATCC and 3’ ACCAGACTTGCCCTCC; and 28S rRNA: 5’ CGGATTGTTTGAGAATGCA and 3’ CAAAGTTCTTTTCAACTTTCCC. The genes were cloned into the pT4P vector for riboprobes and RNA interference food preparation. Three biological repeats were collected from each group for RNA extraction and gene expression analysis.

### Plasmid construction and cell transfection

Each of the full-length FBL homologs in planarians optimized for the human sequence (Genewiz, Inc.) was fused with a 3X Myc tag for transfection to 293T cells, and with 3X Flag tag for transfection to H9 cells. 293T cells were maintained in DMEM (Shanghai Basalmedia Technologies Co., Ltd., L110KJ) with 10% FBS (Excell Bio, FSP500) at 37°C under 5% CO_2_. The 293T cells were seeded in a 6-well plate with coverslips and grown to 70% confluence for transfection with Lipofectamine 3000 reagent (Thermo Fisher Scientific, L3000001) as recommended in the kit literature. H9 cells were cultured in mTeSR™ Plus media (STEMCELL, 100-0276) supplemented with 1% Penicillin-Streptomycin (Gibco, 15140122). 400,000 H9 cells for each condition were electroporated with 1 μg plasmid via a human stem cell nucleofector™ kit (VPH-5012) by Lonza AMAXA Nucleofector 2B.

### Phylogenetic analysis

The protein sequences of the FBL homologs were downloaded from the websites https://www.uniprot.org/ and https://planosphere.stowers.org/. The FASTA files of the sequences were generated by MEGA version 6 (Tamura *et al*, 2013). All the protein sequences were aligned online MAFFT version 7 with the L-INS-i method (https://mafft.cbrc.jp/alignment/server/index.html) (Katoh *et al*, 2019; Kuraku *et al*, 2013) and trimmed in MEGA 6. Maximum likelihood analyses were run using the online web server IQ TREE (http://iqtree.cibiv.univie.ac.at/) (Hoang *et al*, 2018; Nguyen *et al*, 2015) with 1,000 ultrafast bootstrap replicates, the WAG amino acid substitution model, four substitution rate categories and the proportion of invariable sites estimated from the dataset. The tree was produced by MEGA 6.

### Antibody staining

After 48 h of transfection, cells on coverslips were fixed with 4% paraformaldehyde (PFA) for 45 min and permeabilized with 0.5% PBSTx. The cells were blocked in 5% horse serum for one hour and incubated with anti-MYC primary antibody (SCBT, 9E10) at a 1:500 dilution or anti-FLAG M2 (Sigma, F1804) at a 1:400 dilution overnight at 4°C. The cells were washed with 0.3% PBSTx four times for 15 min per wash and incubated in Alexa Fluor 488 Goat Anti-Mouse IgG H&L (Abcam, ab150117) at a 1:1000 dilution at room temperature (RT) for two hours. The cells were mounted with ProLong™ Gold Antifade Mountant (Thermo Fisher Scientific, P36934) before being stained with DAPI for 15 min at a 1:1000 dilution.

### RNAi experiment

pT4P with the genes of interest was transferred into the *E. coli* HT115 strain, and a single colony was cultured for 16 h as a starter culture in 2X YT medium. The starter culture was allowed to grow tenfold before induction with 1 mM IPTG for two hours until the OD600 was approximately 0.8. The 4X RNAi food for *fbl-2* KD was prepared by mixing 50 mL of cultured bacteria with 125 μL of liver homogenate (90% liver paste, 5.5% 1X Montjuic water, and 4.5% red food coloring). The 2X concentration of RNAi food was prepared for *fbl-1* KD. The feeding schedule was arranged according to practical necessity (to produce the desired phenotype but no lethality). The *fbl-1* KD and *fbl-2* KD planarians were fed every three days for a total of five and six times, respectively, and collected the samples after 7-day starvation.

### WISH and FISH

The worms were killed with 5% N-acetylcysteine (NAC) for 5 minutes and fixed with 4% formaldehyde (FA) for 45 minutes following reduction and dehydration. After 2 h of incubation in 100% methanol, the worms were used to perform WISH and FISH experiments. The worms were hybridized with riboprobe in Hybe for up to 16 h at 56°C after being bleached in 5% deionized formamide in 0.5X SSC in direct light for 1.5 h and permeabilized in proteinase K for 10 min. For WISH, the worms were incubated with anti-DIG conjugated with AP overnight at 4°C, and the color was developed with NBT and BCIP. For FISH, the worms were incubated with anti-DIG/DNP/FL conjugated with POD or HRP at 4°C overnight and developed with tyramine labeled with fluorescence at RT for one hour. The anti-H3P and anti-6G10 antibody staining was performed after FISH. The anti-Smed-FBL-2 antibody (antigen aa 2-19, Abclonal Inc.) was used at a 1:500 dilution after worms were bleached and permeabilized. The anti-Smed-PIWI-1 antibody was a gift from J. Rink and was used at a 1:10000 dilution.

### Image acquisition and analysis

WISH samples and live worms were imaged on a Leica M205 FA fluorescence stereomicroscope. FISH samples were imaged on a Nikon C2Si inverted confocal microscope. The images were processed in Fiji, Adobe Photoshop, and Adobe Illustrator CC 2018.

## Acknowledgments

We thank all the members of the Lei lab and biomedical research core facilities at Westlake University for their technical support. KL was supported by the National Natural Science Foundation of China (31970750, 32122032) and the start-up fund from the Westlake Education Foundation.

## Author contributions

**Jiajia Chen:** Conceptualization; methodology; validation; investigation; data curation; formal analysis; visualization; writing - original draft; writing - review and editing. **Xue Pan:** Supervision; methodology; validation; investigation. **Hao Xu:** Validation; investigation. **Yuhong Zhang:** Validation; investigation. **Kai Lei:** Conceptualization; methodology; validation; investigation; data curation; formal analysis; visualization; writing - original draft; writing - review and editing.

## Figure legends

**Figure EV1.**
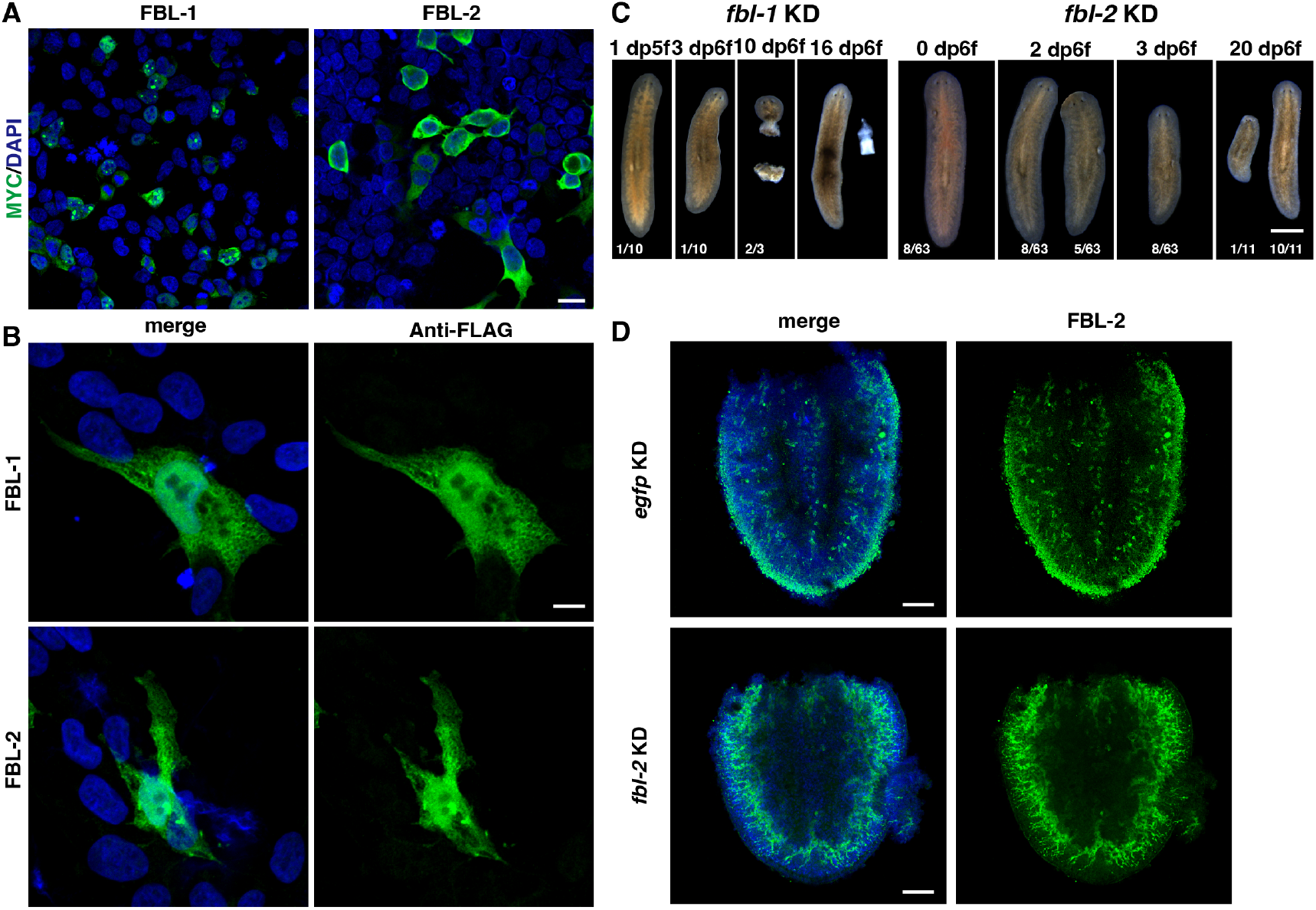
*fbl-1* and *fbl-2* are required for tissue homeostasis. A. Expression analysis of FBL-1 and FBL-2 in 293T cells. Scale bar, 20 μm. B. Expression analysis of FBL-1 and FBL-2 in human embryonic cells H9. Scale bar, 10 μm. C. Animal phenotype observation after KD of *fbl-1* or *fbl-2*. Scale bar, 500 μm. D. Reduced expression level of FBL-2 after *fbl-2* KD at 24 hpa. Scale bars, 100 μm.

**Figure EV2.**
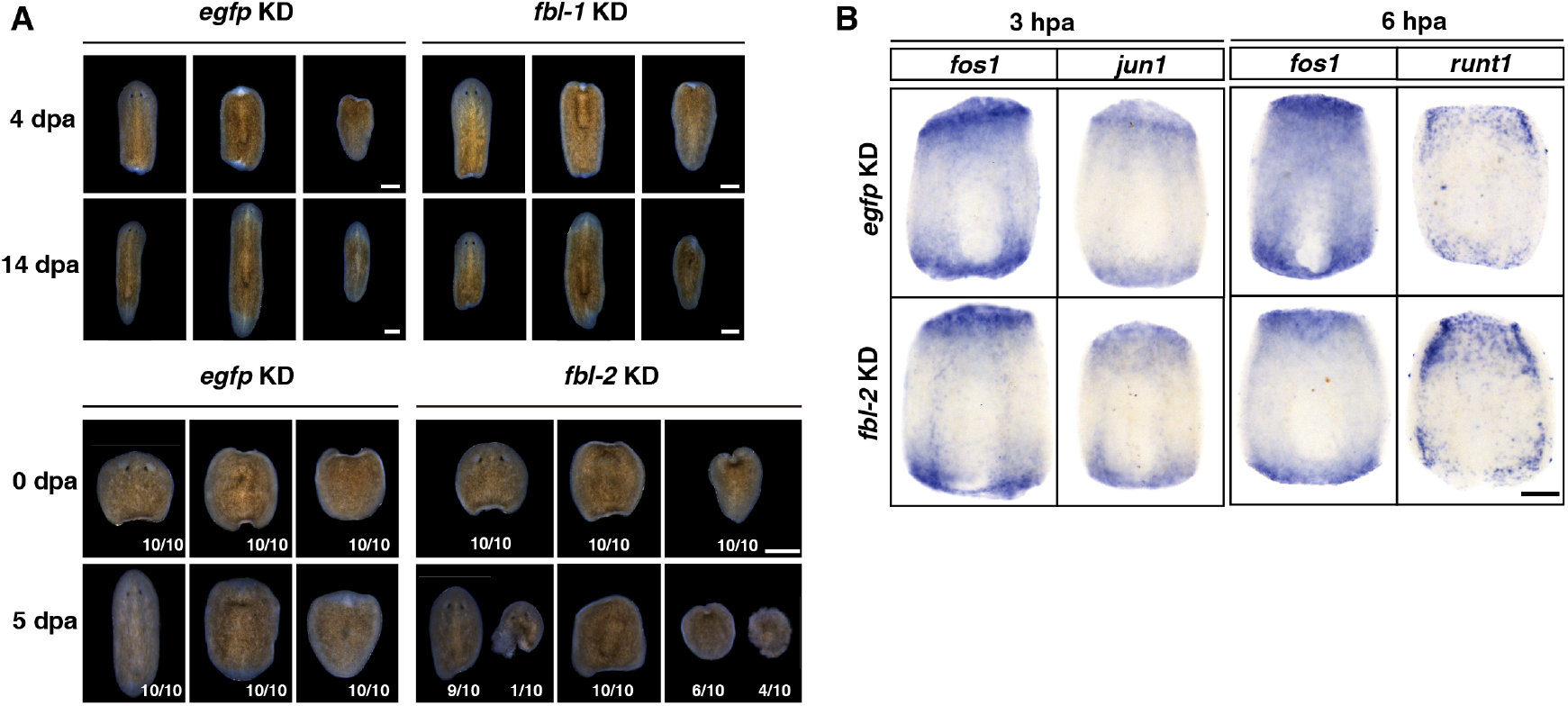
*fbl-1* and *fbl-2* are required for regeneration. A. Defects in regeneration after KD of *fbl-1* at 4, 14 dpa, and KD of *fbl-2* at 0, 5 dpa. Scale bars, 500 μm. B. WISH of wound-induced gene expression levels at 3 and 6 hpa (*jun-1, fos-1, runt1*) in *fbl-2* KD and *egfp* KD animals. Scale bar, 200 μm.

**Figure EV3.**
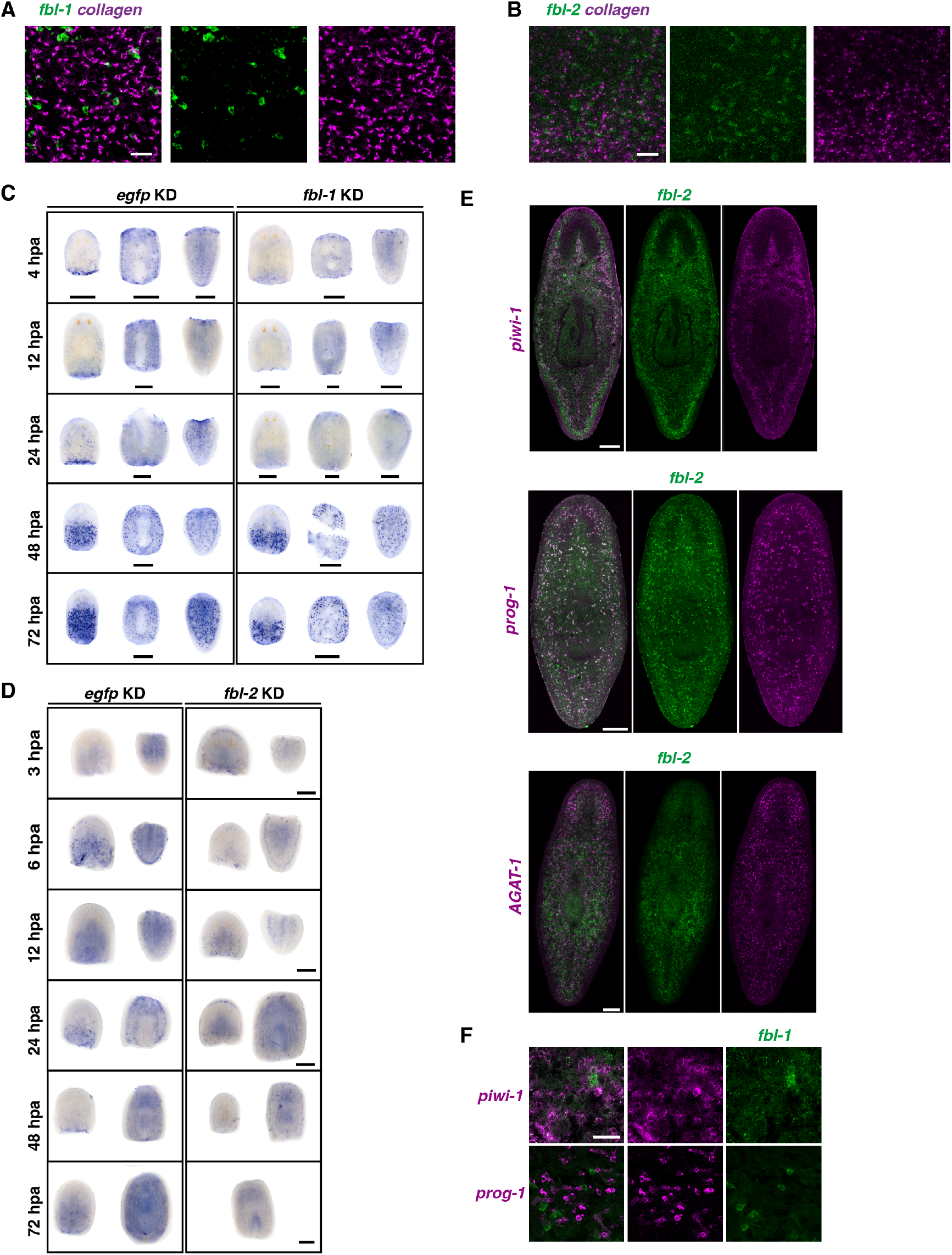
Expression of *fbl-1* and *fbl-2* in distinct cell lineages. A, B. No colocalization of the expression of *fbl-1* and *fbl-2* with collagen. Scale bars, 50 μm. *C. fbl-1* expression at 4, 12, 24, 48, and 72 hpa between the *egfp* KD and *fbl-1* KD groups. Scale bars, 500 μm. *D. fbl-2* expression at 3, 6, 12, 24, 48, and 72 hpa between the *egfp* KD and *fbl-2* KD groups. Scale bars, 500 μm. *E. fbl-2* expression with *piwi-1, prog-1*, and *AGAT-1* in intact animals. Scale bars, 200 μm. *F*. No colocalization of the expression of *fbl-1* with *piwi-1* and *prog-1* at 72 hpa. Scale bar, 100 μm.

**Figure EV4.**
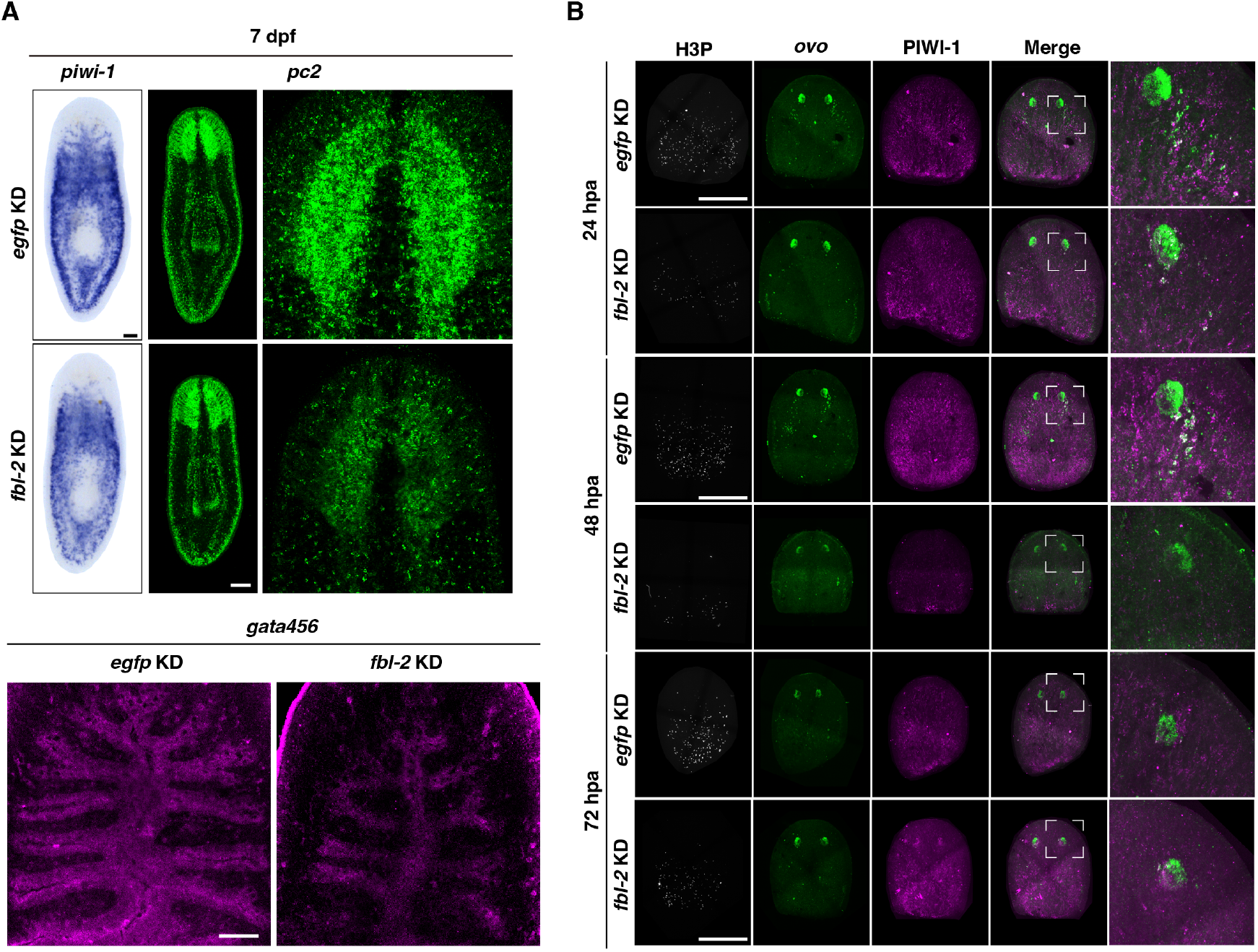
*fbl-2* is a general master regulator of cell differentiation. *A*. Shrinking body size with a slight decrease in the number of stem cells (*piwi-1*) and defects in multiple tissues (neuron, *pc2*; intestine, *gata456*). Scale bars, 100 μm. *B. fbl-2* KD blocks *ovo*^+^ cell differentiation at 24, 48, and 72 hpa. Scale bars, 200 μm.

**Figure EV5.**
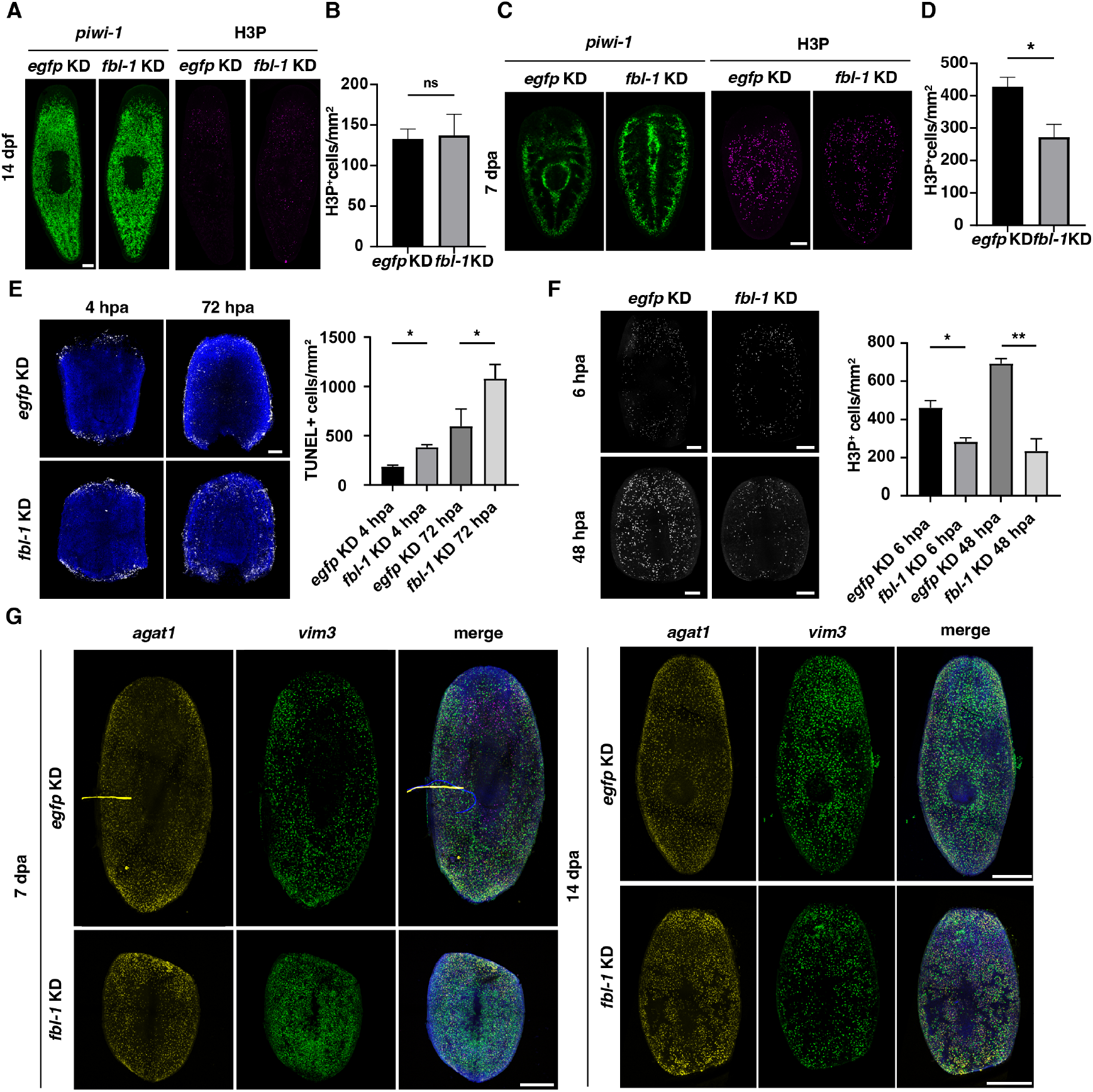
*fbl-1* inhibition causes secondary defects in stem cell behaviors. *A*. No significant changes in stem cell numbers or proliferative cells during homeostasis after *fbl-1* KD. Scale bar, 200 μm. *B*. Quantification of H3P^+^ cells at 14 dpf after *fbl-1* KD. *C*. Cell proliferation decreased, but stem cell differentiation increased during regeneration in *fbl-1* KD. Scale bar, 200 μm. *D*. Quantification of H3P^+^ cells at 7 dpa after *fbl-1* KD compared with those in *egfp* KD animals. \**P* < 0.05 (Student’s *t*-test). *E*. TUNEL staining shows Increased cell death in *fbl-1* KD animals at 4 hpa and 72 hpa. Scale bar, 200 μm. *F*. Reduced cell proliferation was revealed by fluorescence staining and quantification of the H3P^+^ cells in *fbl-1* KD animals at 6 hpa and 48 hpa. Scale bar, 200 μm. \**P* < 0.05; \*\**P* < 0.01 (Student’s *t*-test). *G*. Expression of *AGAT-1*^+^ and *vim3*^+^ cells during regeneration at 7 dpa and 14 dpa in *egfp* KD and *fbl-1* KD animals. Scale bars, 500 μm.

